# A theoretical framework for aerodynamic braking and landing in gliding mammals

**DOI:** 10.64898/2026.07.27.740869

**Authors:** Kota Nojiri

## Abstract

Gliding enables mammals to forage and escape from predators by moving between discontinuous forests. Its benefits depend not only on glide distance but also on the ability to decelerate and land safely. This study developed a theoretical framework linking glide distance, gliding velocity, aerodynamic braking, body mass, and braking distance. Twenty-two representative distance-velocity observations from eight studies and five species were compiled. Among distance-velocity models, log-distance and saturated with *V*_0_ models received nearly equivalent support. Both models predicted increasing velocity with glide distance, with the rate of increase declining at longer distances. For a 1 kg animal undergoing a 60% reduction in velocity, predicted kinetic energy remaining immediately before contact increased from 4.80-5.14J at 20 m to 8.13-8.90 J at 80 m. This velocity reduction corresponded to a dissipation of 84% of approach kinetic energy before contact. Over a braking distance of 1 m, the required mean deceleration increased from 2.57-2.75 *g* at 20 m to 4.35-4.77 *g* at 80 m. At 80 m, shortening the braking distance from 4 to 0.5 m increased the required deceleration from 1.09-1.19 to 8.70-9.53 *g*. These results indicate that the absolute energetic and deceleration requirements of landing increase with glide distance, even when velocity is close to an asymptote. These results provide a quantitative basis for considering aerodynamic braking and landing requirements alongside conventional measures of glide performance.

**Summary statement:** Models quantify how glide distance, aerodynamic braking, and braking distance affect pre-contact kinetic energy and deceleration requirements in gliding mammals.

## Introduction

Gliding has evolved independently in numerous animal lineages and occurs in mammals, reptiles, amphibians, fishes, and arthropods (Dudley et al., 2007; Socha et al., 2015). In mammals, gliding enables animals to cross discontinuities in the forest canopy without descending to the ground and has been proposed to reduce travel time, increase access to spatially distributed food resources, and facilitate predator avoidance (Byrnes and Spence, 2011; Dial, 2003; Dudley et al., 2007). Gliding has also frequently been assumed to reduce the energetic cost of arboreal locomotion by replacing horizontal quadrupedal movement with aerial travel, though empirical studies indicate that its energetic benefits depend on body size and environmental context, and are not universal among species (Byrnes et al., 2011; Flaherty et al., 2010). Thus, the selective advantages of gliding may involve a combination of travel speed, access to resources, predator avoidance, and energetic economy rather than the maximization of glide distance alone.

The benefits of gliding, nevertheless, depend on the animal’s ability to terminate each glide safely. As an animal accelerates during aerial descent, generated kinetic energy must ultimately be dissipated either aerodynamically before landing or mechanically at contact. The translational kinetic energy immediately before contact is proportional to body mass and to the square of landing velocity. Consequently, relatively small increases in velocity may substantially increase the energy that must be dissipated before and during landing. Landing performance may therefore impose an important mechanical constraint on the evolution of gliding distance, velocity, and body size.

Gliding mammals can actively modify the aerodynamic forces acting on the body by changing limb position, body pitch, angle of attack, and patagial configuration (Bishop, 2007, 2006; Byrnes et al., 2011; Socha et al., 2015). Before landing, they present a larger surface area against the direction of travel by pitching the body upwards and extending their patagium, thereby increasing drag and reducing velocity before contact (Byrnes et al., 2008; Paskins et al., 2007). The patagium should therefore be considered not only as a lift-generating surface but also as a structure involved in braking and landing control.

Despite the potential importance of braking and landing to interpret gliding locomotion, quantitative comparisons remain difficult because many gliding mammals are small and nocturnal, making the complete gliding and landing sequence difficult to record in the wild (Goldingay and Scheibe, 2000). In particular, it remains unclear how the relationship between glide distance and velocity affects the kinetic energy remaining at landing and the deceleration required to attain a given landing velocity.

In this study, we compiled published measurements of glide distance and velocity from five species of gliding mammals and developed a theoretical framework linking glide distance, predicted gliding velocity, aerodynamic velocity reduction, body mass, and braking distance. First, linear, log-distance, and asymptotic models representing alternative distance-velocity relationships were compared. Then, we used the supported models to estimate the kinetic energy remaining immediately before contact and the mean constant deceleration required to achieve specified proportional reductions in velocity over different braking distances. This framework does not estimate contact force directly but provides a basis for evaluating how braking and landing requirements may constrain the evolution of gliding performance in mammals.

## Materials and Methods

All data processing, model fitting, and graphical analyses were conducted in R (version 4.4.1) (R Core Team, 2024). AICc values were calculated using the MuMIn package (version 1.48.19) (Bartoń, 2026).

### a Compilation of published glide-performance data

Published data on glide distance, gliding velocity, and body mass were compiled for gliding mammals. The final dataset contained 22 representative distance-velocity data points obtained from eight published studies and included five species: *Acrobates pygmaeus*, *Galeopterus variegatus*, *Glaucomys sabrinus*, *Petaurista leucogenys*, and *Petaurista petaurista* (Table 1).

**Table 1.**
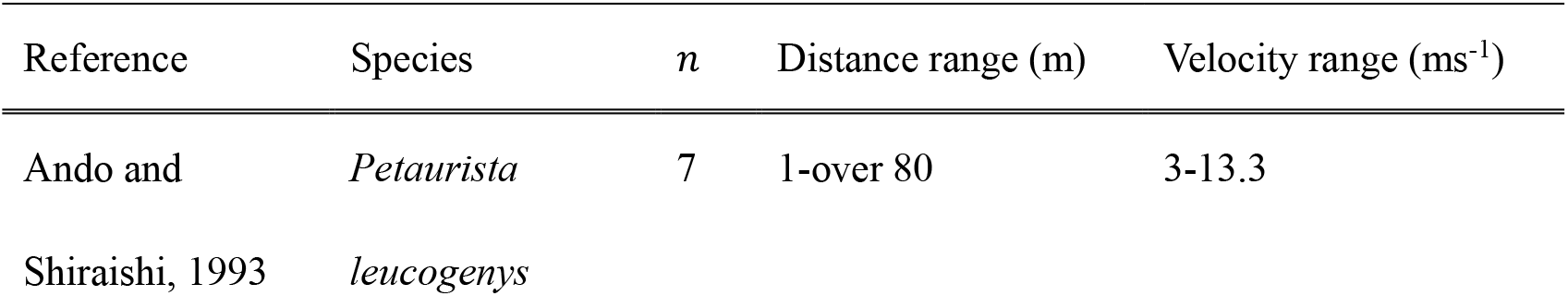
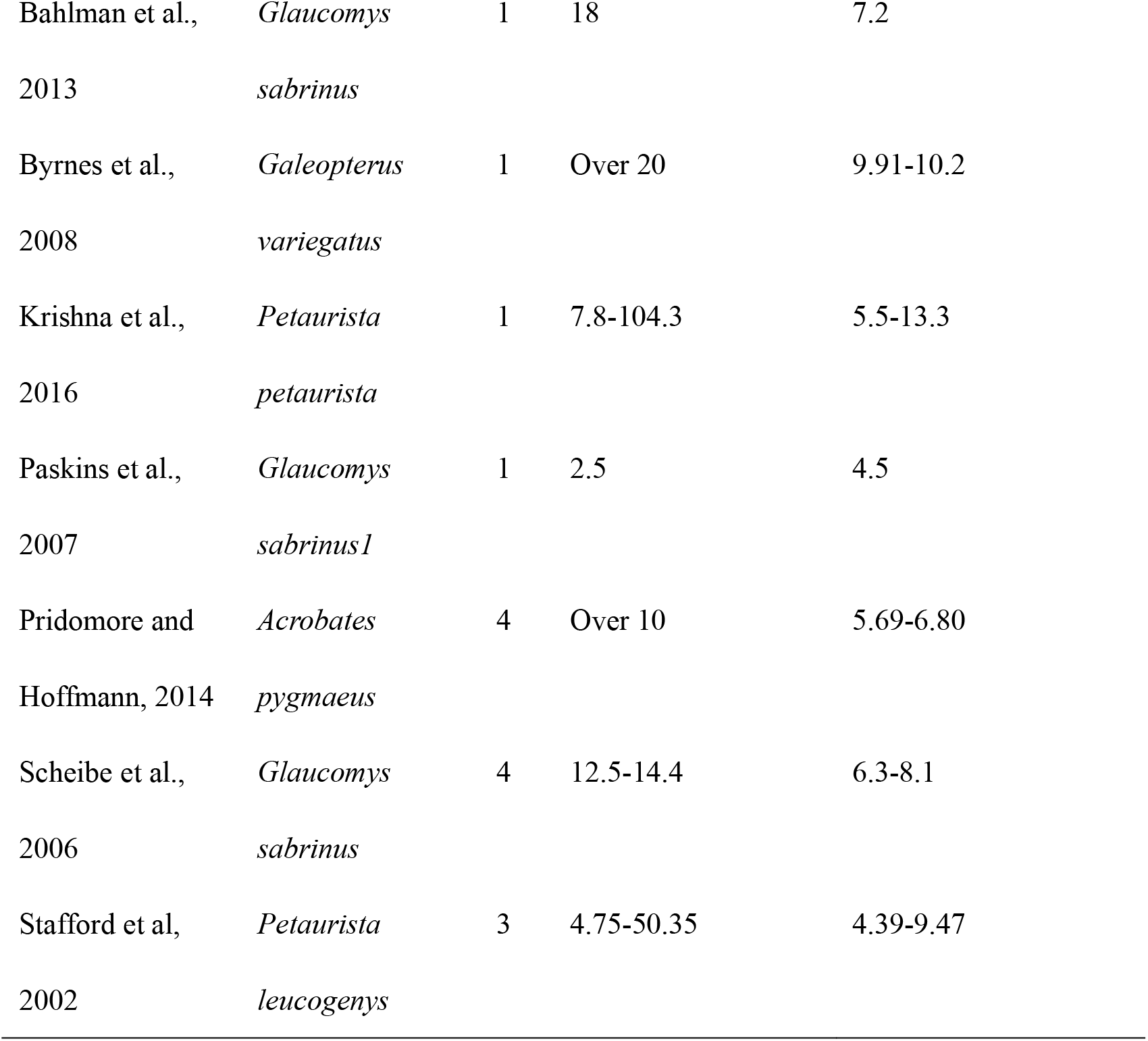
Published glide-performance data used to model the relationship between glide distance and velocity. *n* is the number of representative distance-velocity observations included from each study. Distance and velocity ranges indicate the minimum and maximum representative values included in the analysis. Complete data are provided in Table S1.

Data were extracted according to the form in which glide performance was reported in each study. When mean glide distance and mean velocity were reported, these values were used directly. When glide distance and velocity were reported as ranges, the reported minimum distance was paired with the reported minimum velocity, and the reported maximum distance was paired with the reported maximum velocity. An additional representative point was generated using the arithmetic midpoint of each reported range. When only a lower bound for glide distance was reported (e.g., “over 80m”), the reported threshold distance was treated as the representative minimum distance and paired with the reported or calculated mean glide velocity.

Body mass values reported in the original studies were used whenever available. When an exact body mass was not provided, the species-specific body mass ranges reported in other literature were used (Nowak, 1999a, 1999b), and the mean of the reported range was adopted as the representative body mass. Representative body mass, glide distance, and glide velocity were denoted by *m*, *D*, and *V*, respectively.

### b Distance-velocity models

Gliding velocity was modeled as a function of glide distance. Four candidate models were fitted to represent alternative relationships between distance and velocity.

The linear model assumed that velocity increased at a constant rate with glide distance:

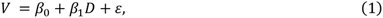

where *β*_0_ is the intercept, *β*_1_ is the slope, and *ε* is the residual error.

The log-distance model assumed that velocity increased rapidly over short glide distances, but that rate of increase declined as distance increased.

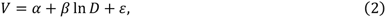

where *α* is the intercept and *β* describes the change in velocity with the natural logarithm of glide distance.

The first saturated model assumed that the glide velocity approached an asymptotic maximum from a velocity of zero at *D* = 0.

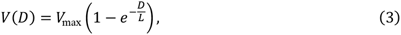

where *V*_max_ is the asymptotic maximum velocity, and *L* is the characteristic distance controlling the rate at which the asymptote is approached.

The second saturated model included a non-zero velocity parameter at *D* = 0.

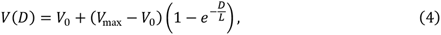

where *V*_0_ is the velocity predicted at zero glide distance.

The linear and log-distance models were fitted by ordinary least-squares regression. The two saturated models were fitted by nonlinear least-squares regression. For the first saturated model, starting values were set to 1.2 times the maximum observed velocity for *V*_max_ and the median observed glide distance for *L*. For the second saturated model, starting values were set to the minimum observed velocity for *V*_0_, 1.2 times the maximum observed velocity for *V*_max_, and the median observed distance for *L*. These parameters were constrained to positive values.

### c Model comparison and prediction

The four candidate models were compared using the Akaike information criterion corrected (AICc) for small sample size. For each model, the difference from the minimum AICc was calculated as

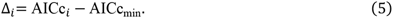

The relative likelihood of each model was calculated as

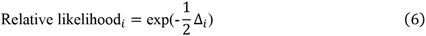

and the corresponding AICc weight was calculated by dividing each relative likelihood by the sum of the relative likelihoods across the models. Residuals were calculated as the observed velocity minus the fitted velocity.

Residual patterns were examined against fitted values and glide distance to identify systematic deviations from the fitted relationships. The log-distance model and the saturated with *V*_0_ model were retained as alternative candidate descriptions of the distance-velocity relationship. Both models were used in the subsequent theoretical calculations. Predicted gliding velocities were calculated at 500 equally spaced distances spanning the minimum to maximum glide distances represented in the dataset. Predictions were therefore restricted to the observed distance range and were not extrapolated beyond the available data.

For the subsequent theoretical calculations, the velocity predicted from each model was treated as the representative velocity immediately before aerodynamic braking. This velocity was denoted by

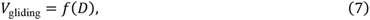

where *f*(*D*) represents either the fitted log-distance model or the fitted saturated with *V*_0_ model.

This treatment assumes that the representative gliding velocities compiled from the literature provide an approximation of pre-braking velocity. The conceptual relationship among glide distance (*D*), braking distance (*S*_brake_), gliding velocity (*V*_gliding_), and landing velocity (*V*_landing_) is illustrated in Fig. 1. The overall workflow of the study is summarized in Fig. 2.

**Figure 1.**
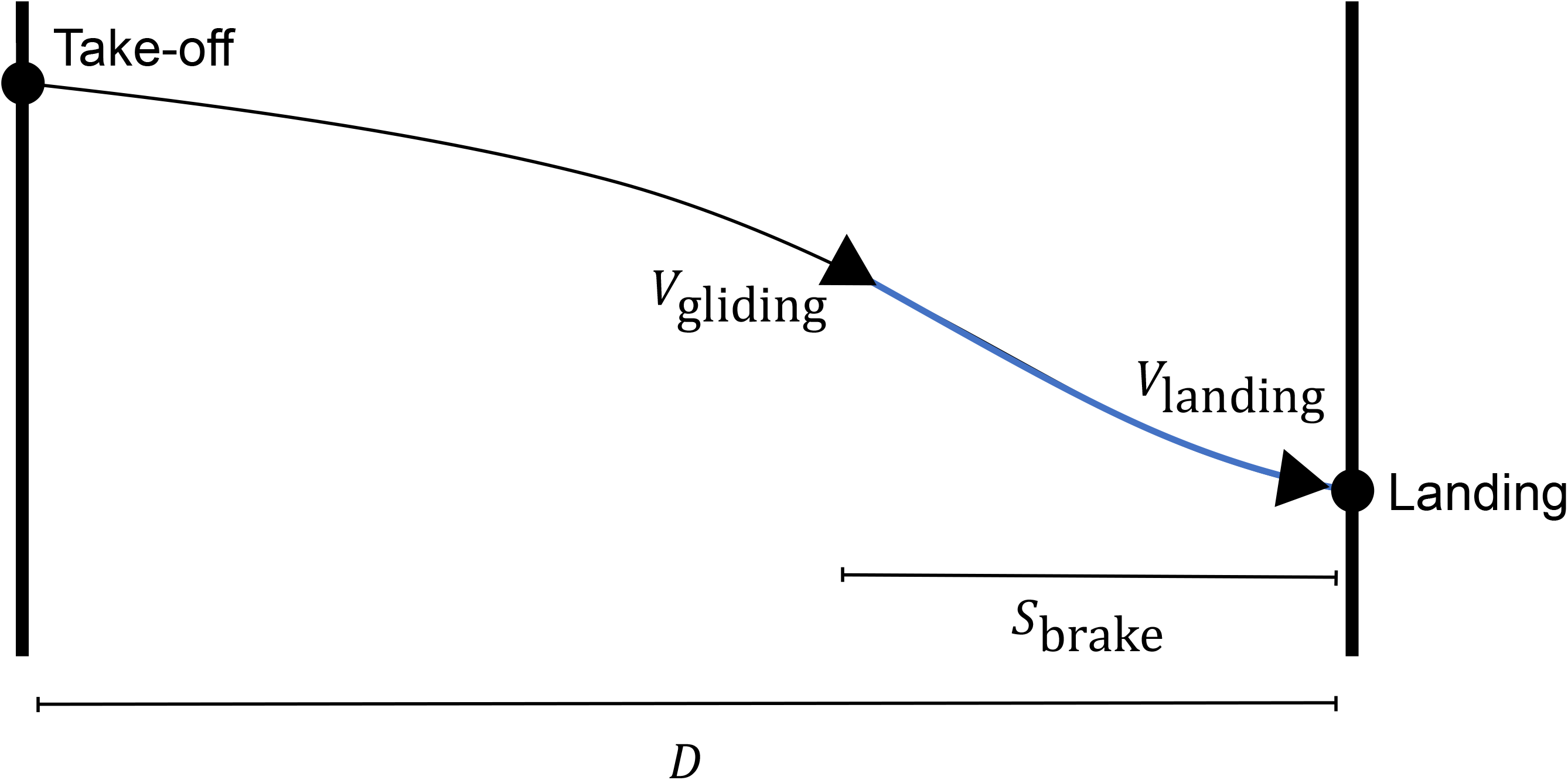
Conceptual framework of the gliding and aerodynamic braking model. Black line represents glide trajectory, and blue line indicates the aerodynamic braking phase. The animal glides over a horizontal distance (*D*) before aerodynamic braking reduces the velocity from *V*_gliding_ to *V*_landing_ over the braking distance (*S*_brake_).

**Figure 2.**
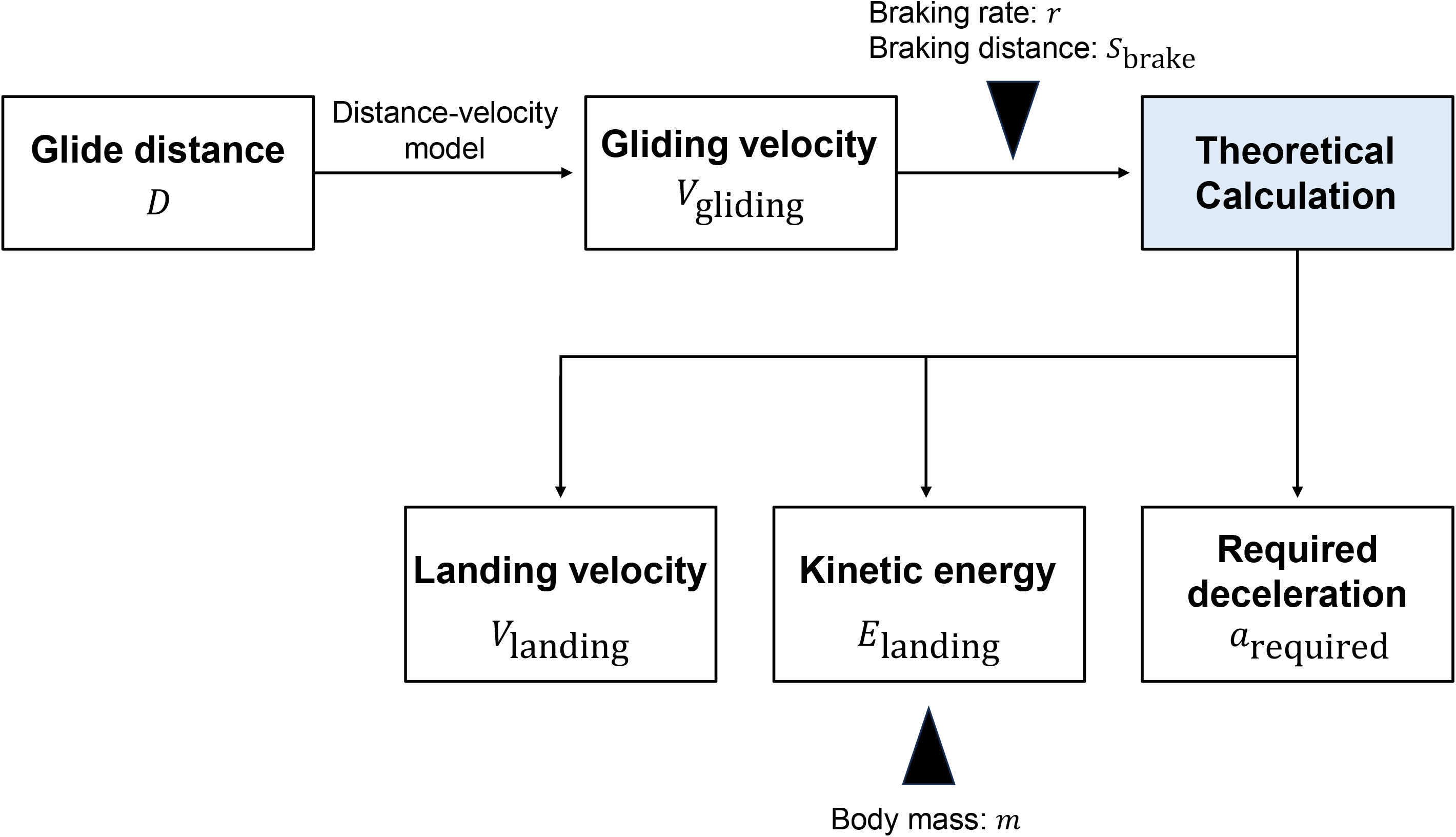
Workflow of the theoretical calculations. Glide distance (*D*) was first converted to gliding velocity (*V*_gliding_) using the fitted distance-velocity models. Theoretical calculations were then performed using the braking rate (*r*) and braking distance (*S*_brake_). Predicted landing velocity (*V*_landing_), landing kinetic energy (*E*_landing_), and the required aerodynamic deceleration (*a*_required_) were subsequently calculated. Body mass (*m*) was incorporated only in the kinetic energy calculations.

### d Aerodynamic braking and landing velocity

Landing velocity was modelled as a fixed proportional reduction in the glide velocity predicted immediately before braking:

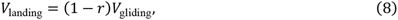

where *r* is the proportional reduction in velocity during aerodynamic braking.

Velocity-reduction ratio scenarios of *r* = 0.40, 0.50, and 0.60 were evaluated to examine the sensitivity of the theoretical predictions to braking performance. A 60% reduction in velocity was used as the principal scenario for numerical summaries (Byrnes et al., 2008). Under this scenario, the landing velocity was 40% of the predicted gliding velocity.

### e Partitioning of kinetic energy

The kinetic energy immediately before braking (*E*_approach_) was partitioned into energy dissipated aerodynamically before contact (*E*_air_) and kinetic energy remaining at landing (*E*_landing_):

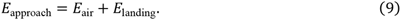

*E*_approach_ was calculated as

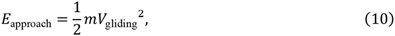

and *E*_landing_ was calculated as

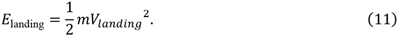

Here, *E*_landing_ represents the translational kinetic energy that remains to be dissipated after contact through interactions among the limbs, musculoskeletal system, whole-body motion, and the landing substrate. It does not directly represent impact force or tissue stress. *E*_air_ was calculated as

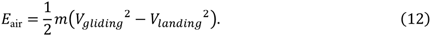

The proportion of approach kinetic energy dissipated before landing (*φ*_air_) was calculated as

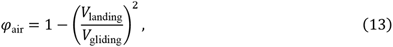

Substitution of Eqn. 8 gives

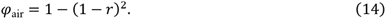

Kinetic energy calculations were conducted for hypothetical body masses of 0.1, 0.5, 1.0, 1.5, and 2.0 kg. This range was used to illustrate how absolute landing energy scales with body mass while retaining the same predicted distance-velocity relationship and braking performance.

Numerical summaries were generated principally for a body mass of 1 kg and a 60% reduction in velocity (Byrnes et al., 2008) at glide distances of approximately 20, 40, 60, and 80 m.

### f Required aerodynamic deceleration

The magnitude of the constant deceleration required to reduce *V*_gliding_ to *V*_landing_ over a prescribed braking distance was estimated from the constant-acceleration kinematic relationship

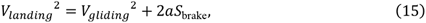

where *a* is acceleration along the direction of motion, and *S*_brake_ is braking distance. Because acceleration is negative during braking, the positive magnitude of the required deceleration was calculated as

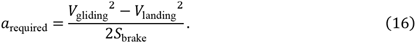

Required deceleration was normalized by gravitational acceleration and expressed in units of *g* as

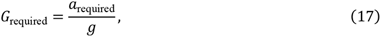

where *g* = 9.81 m s^-2^.

Braking distances of 0.5, 1.0, 2.0, and 4.0 m were evaluated in combination with velocity reductions of 40, 50, and 60% (Byrnes et al., 2011; Paskins et al., 2007). Required deceleration was summarized across glide distances for a braking distance of 1 m and across braking distances for a glide distance of approximately 80 m. The estimated values represent the mean constant aerodynamic deceleration required over the prescribed braking distance. They do not represent peak external forces experienced at contact or forces transmitted directly through individual limbs.

## Results

### a Relationship between glide distance and velocity

The log-distance and saturated with *V*_0_ models received nearly equivalent support based on AICc (Table 2). The log-distance model had the lowest AICc (AICc = 70.88, AICc weight = 0.51), followed closely by the saturated with *V*_0_ model (AICc = 71.07, ΔAICc = 0.19, AICc weight = 0.46). By contrast, the linear model (AICc = 76.44, ΔAICc = 5.56, AICc weight = 0.03) and the saturated model constrained to pass through the origin (AICc = 90.76, ΔAICc = 19.88, AICc weight < 0.001) received little support. Based on the results, these two models were adopted for further prediction.

**Table 2.** Comparison of candidate models describing the relationship between glide distance and velocity. ΔAICc is the difference between the AICc of each model and the minimum AICc (Eqn. 5). Relative likelihood was calculated as Eqn.6. Models are arranged in ascending order of AICc.

| Model | $AICc$ | $\Delta AICc$ | Relative likelihood | $AICc$ weight |
| --- | --- | --- | --- | --- |
| Log-distance | 70.88 | 0 | 1 | 0.51 |
| Saturated with $V_0$ | 71.07 | 0.19 | 0.91 | 0.46 |
| Linear | 76.44 | 5.56 | 0.06 | 0.03 |
| Saturated through origin | 90.76 | 19.88 | < 0.0001 | < 0.001 |

The log-distance model (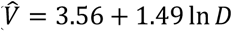, *F*_1,20_ = 79,76, *p* = 2.04 × 10^-8^, Adjusted R^2^ = 0.79) and the saturated with *V*_0_ model 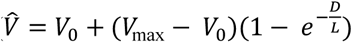, *V*_max_ = 10.88, *L* = 26.63 m; residual SE = 1.04) both described a positive relationship between glide distance and velocity, but the rate of increase declined at longer distances (Fig. 3). The models predicted a rapid increase in velocity over short glide distances, followed by a progressively slower increase at longer distances. Parameter estimates for all candidate models are provided in Table S2, and their fitted relationships and residual patterns are shown in Figs. S1 and S2, respectively.

**Figure 3.**
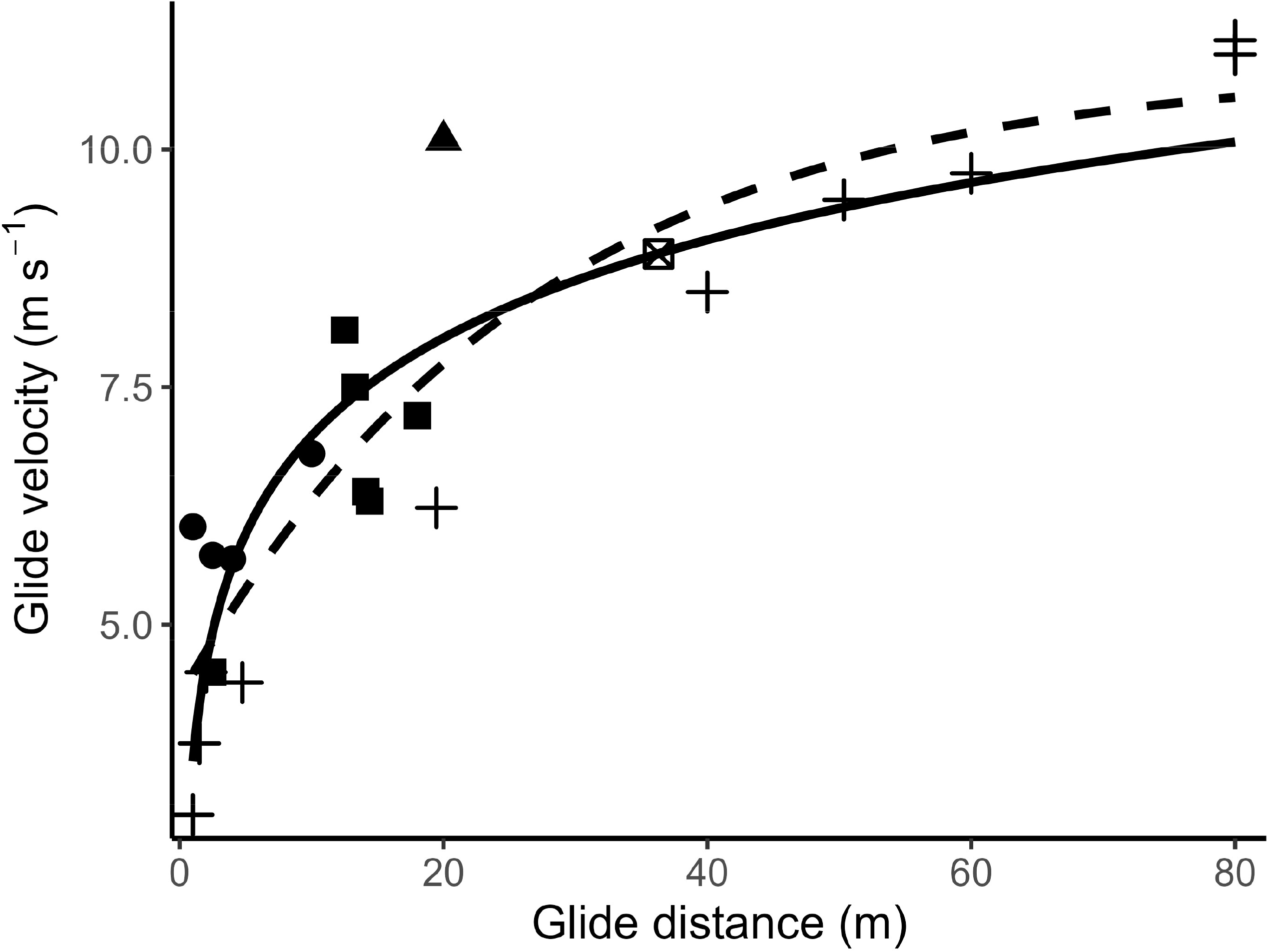
Relationship between glide distance and gliding velocity in gliding mammals. Points represent velocity estimates compiled from the literature (circles, *Acrobates pygmaeus*; triangles, *Galeopterus variegatus*; squares, *Glaucomys sabrinus*; crosses, *Petaurista leucogenys*; and crossed squares, *Petaurista petaurista*). Lines show predictions from the two best-supported distance-velocity models: the log-distance model (solid) and the saturated with *V*_0_ model (dashed).

### b Effects of aerodynamic braking on landing velocity and energy

Predicted landing velocity increased with glide distance under all braking scenarios, reflecting the distance-dependent increase in predicted gliding velocity (Fig. 4A). Greater proportional reductions in velocity substantially decreased the velocity remaining immediately before contact.

**Figure 4.**
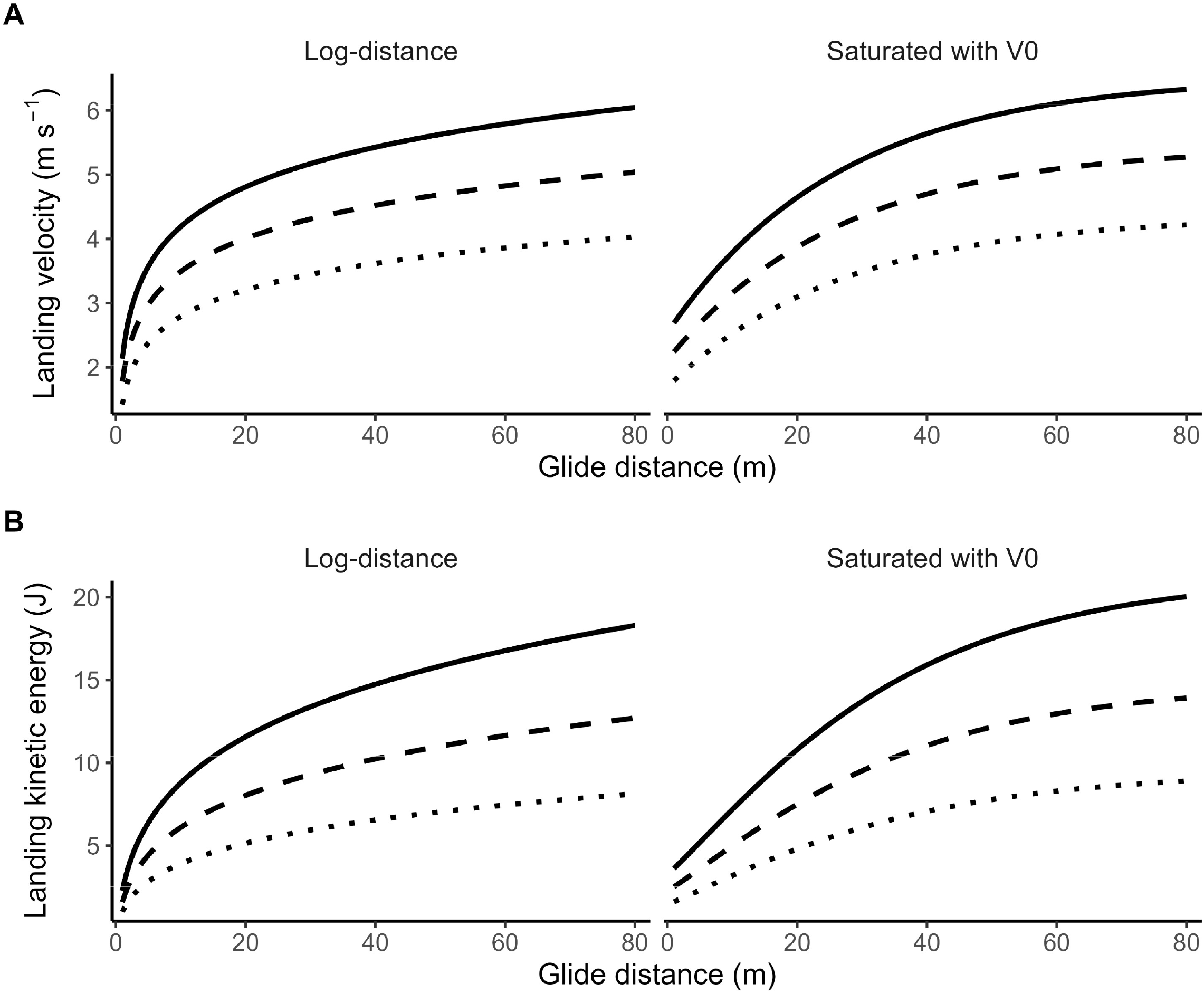
Predicted effects of aerodynamic braking on landing velocity and kinetic energy across glide distances. (A) Predicted landing velocity following reductions in gliding velocity of 40 (solid), 50 (dashed), and 60% (dotted). (B) Corresponding kinetic energy remaining immediately before landing for a body mass of 1 kg. Predictions are shown separately for the log-distance and saturated with *V*_0_ models.

Predicted landing kinetic energy increased with glide distance under both the log-distance and saturated with *V*_0_ models (Fig. 4B). For a body mass of 1 kg with a 60% reduction in glide velocity, landing kinetic energy increased from 5.14 to 8.13 J under the log-distance model and from 4.80 to 8.90 J under the saturated with *V*_0_ model between glide distances of 20 and 80 m (Table S3).

The proportion of approach kinetic energy dissipated before contact depended on the proportional reduction in velocity (Eqn. 14). Reductions in velocity of 40, 50 and 60% corresponded to dissipation of 64, 75 and 84% of the approach kinetic energy, respectively. Thus, under the principal 60% velocity-reduction scenario, 84% of the approach kinetic energy was dissipated before contact and 16% remained as landing kinetic energy, irrespective of body mass or glide distance (Fig. 5). Nevertheless, because approach kinetic energy increased with glide distance, the absolute amounts of both energy dissipated before contact and energy remaining at landing also increased with glide distance. Absolute landing kinetic energy increased proportionally with body mass across the examined mass range (0.1, 0.5, 1.0, 1.5, and 2.0 kg) (Fig. S3).

**Figure 5.**
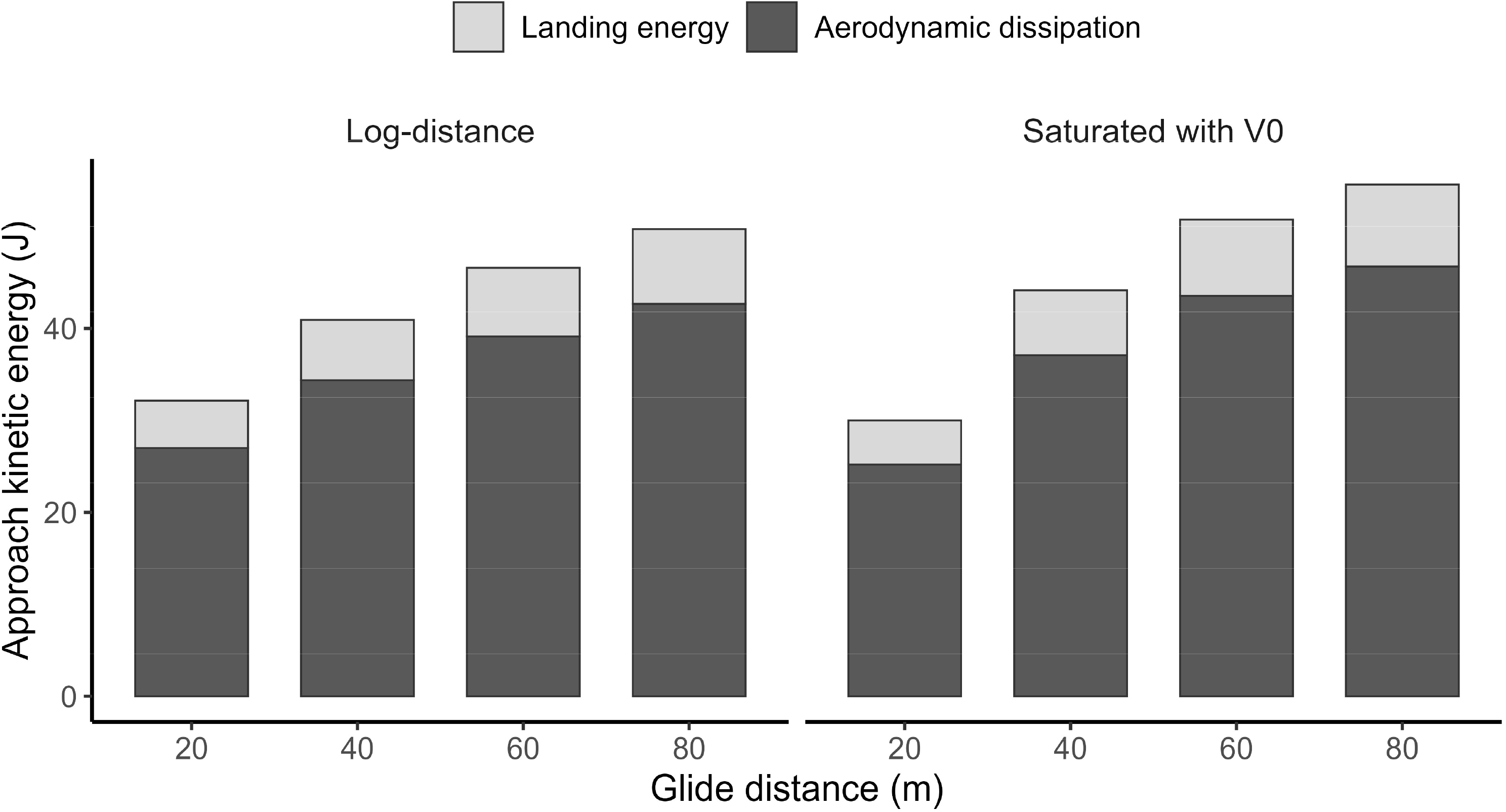
Partitioning of approach kinetic energy during aerodynamic braking across glide distances. Bars show the partitioning of kinetic energy immediately before braking into energy dissipated before contact through aerodynamic braking and kinetic energy remaining immediately before landing. Calculations assume a body mass of 1 kg and a 60% reduction in gliding velocity. Predictions are shown separately for the log-distance and saturated-with *V*_0_ models.

### c Deceleration required before landing

The deceleration required to attain a given proportional reduction in velocity increased with glide distance and with decreasing braking distance (Fig. 6). For a braking distance of 1 m and a 60% reduction in velocity, the required deceleration predicted by the log-distance model increased from 2.75 *g* at a glide distance of 20 m to 4.35 *g* at 80 m. The saturated with *V*_0_ model produced corresponding estimates ranging from 2.57 to 4.77 *g*.

**Figure 6.**
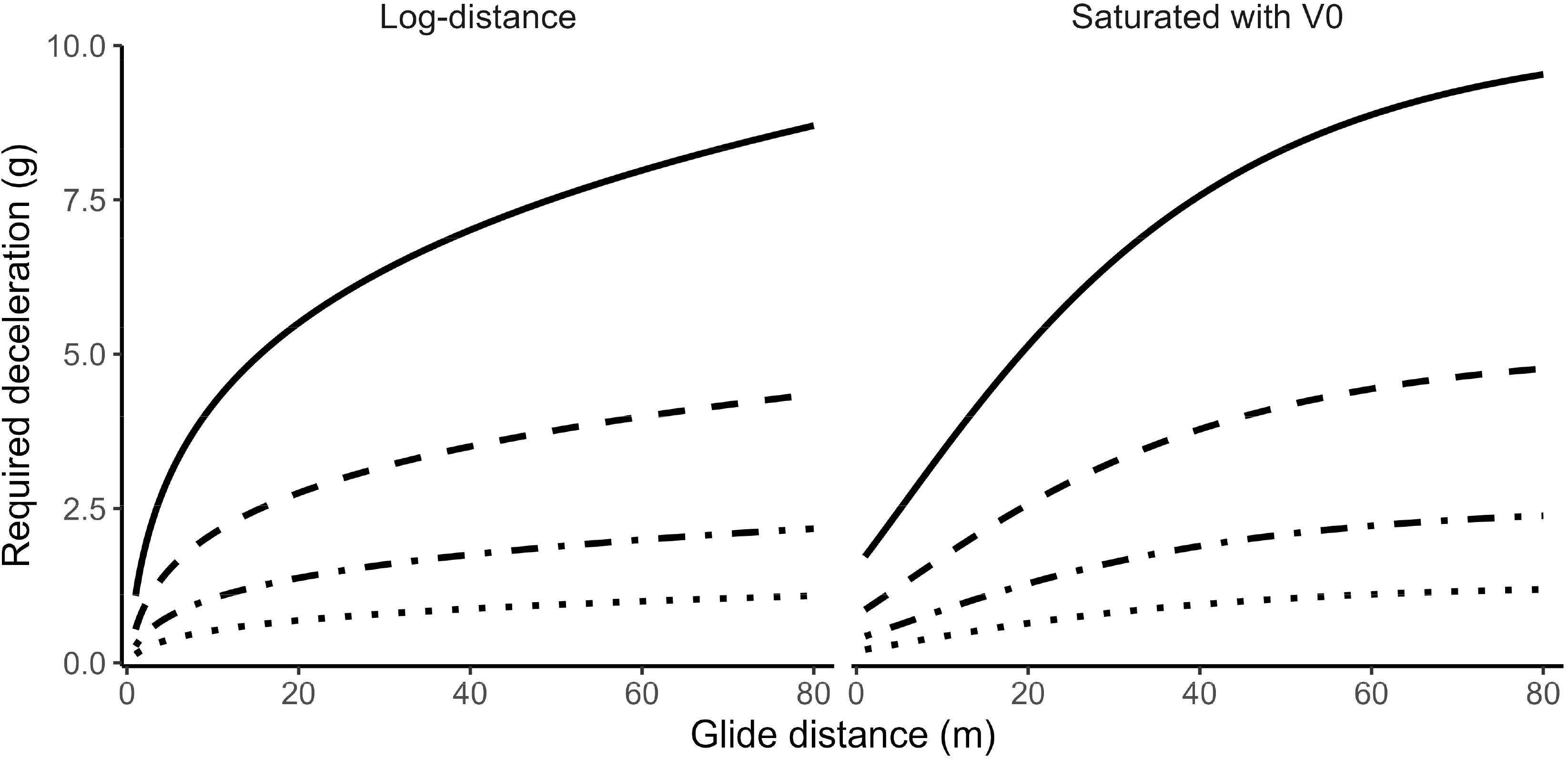
Required deceleration for aerodynamic braking across gliding distances. Required deceleration was calculated assuming a 60% reduction in gliding velocity before landing. Lines represent braking distances of 0.5 (solid), 1.0 (dashed), 2.0 (dot-dashed), and 4.0 m (dotted). Predictions are shown separately for the log-distance and saturated with *V*_0_ models.

At a glide distance of 80 m, shortening the braking distance from 4 to 0.5 m increased the required deceleration from approximately 1.09-1.19 to 8.70-9.53 *g*, depending on the distance-velocity models. The effects of the assumed velocity reduction on predicted landing variables and required deceleration are summarized in Table S4 and Fig. S4.

## Discussion

### a Distance-dependent energetic consequences of gliding

Gliding has generally been interpreted as an adaptation that facilitates predator avoidance, expands the area available for foraging, and reduces the energetic costs associated with moving across the forest (Dudley et al., 2007). However, these benefits depend not only on the capacity to generate lift and travel between distant trees, but also on the capacity to dissipate kinetic energy and control the body before landing (Paskins et al., 2007). The present models provided a quantitative framework for examining this less-considered component of gliding performance.

Both the log-distance and saturated with *V*_0_ models indicated that gliding velocity increases with glide distance, although the rate of increase declined at longer distances (Fig. 3). The similarity of the predictions from these two models suggests that this general tendency was not strongly dependent on whether velocity was assumed to continue increasing logarithmically or to approach an asymptotic maximum. Nevertheless, the models imply different behavior beyond the observed range, and the present results should not be extrapolated to extremely short or long glides.

Although the rate of velocity increase declined with glide distance, the absolute kinetic energy remaining before contact increased with glide distance (Fig. 4B). Thus, a declining marginal increase in velocity does not eliminate the mechanical consequences of longer glides. Because kinetic energy depends on the square of velocity, even relatively small increases in velocity can increase the amount of energy that must be dissipated aerodynamically before contact or absorbed during landing (Paskins et al., 2007). These results suggest that the benefits of increasing glide distances may ultimately be constrained by the animal’s capacity for aerodynamic braking, postural control and energy absorption during landing.

### b Braking distance and landing mechanisms

The required deceleration increased as the prescribed braking distance decreased (Fig. 6). This result emphasizes that landing performance depends not only on approach velocity but also on the distance available for braking. These results highlight the importance of aerodynamic braking as a mechanism for dissipating kinetic energy before contact and thereby reducing the mechanical demands of landing. Consequently, short aerial movements may provide insufficient distance for animals to establish an effective gliding posture and reduce velocity before contact, even when the absolute approach velocity is relatively low.

In northern flying squirrels (*Glaucomys sabrinus*), landing force increased with horizontal range over the relatively short distances examined experimentally, and steep approaches restricted the animal’s ability to pitch upwards and distribute the landing force across all four limbs (Paskins et al., 2007). In contrast, landing forces measured in free-ranging Malayan colugos (*Galeopterus variegatus*) were negatively associated with glide distance, despite greater propulsive impulses during longer glides (Byrnes et al., 2008). The approach of glide velocity towards an asymptotic or terminal value may partly explain why the landing force does not continue to increase at longer distances. However, velocity saturation alone would predict a plateau rather than a decline in landing force. This empirical relationship is not generated directly by the present models, because braking performance was specified as a fixed proportional reduction in velocity. Instead, it indicates that braking rate, braking distance, or landing posture may change with glide distance in living animals. Incorporating such behavioral parameters would be necessary for a model to reproduce a decline in landing force with increasing glide distance.

### c The patagium as a braking system

According to Eqn. 14, under the 60% velocity-reduction scenario, only 16% of the approach energy remained to be dissipated during or after contact, irrespective of body mass or glide distance (Fig. 5). This value should not be interpreted as a direct measurement of energy dissipated by the patagium. Rather, it quantifies the aerodynamic energy dissipation required if animals achieve the velocity reductions reported in previous studies (Byrnes et al., 2011; Paskins et al., 2007). The calculation illustrates that a large proportion of the approach kinetic energy must be removed before contact for the landing velocity to be reduced substantially.

As behavioral observations suggest, the patagium functions not only as a lift-generating surface but also as a structure for producing drag and controlling body orientation (Bahlman et al., 2013; Bishop, 2006, 2007; Pridmore and Hoffmann, 2014). Before contact, gliding mammals increase body pitch and orient the patagium more nearly perpendicular to the direction of trajectory. This behavior enables rapid deceleration and allows the remaining landing force to be distributed across the four limbs.

The low aspect ratio of the patagium may also reflect requirements that extend beyond maximizing glide efficiency (Bishop, 2007), because high-aspect-ratio wings generally reduce induced drag and increase lift-to-drag ratio (Bramesfeld, 2010; Traub, 2013). This raises the possibility that patagial shape may reflect a compromise between glide efficiency and braking performance, although testing this hypothesis will require explicit incorporation of patagial morphology into future models.

### d Limitations and future directions

The present models were constructed from a limited compilation of observed glide distances and velocities in the literature. Direct measurements of complete glide and landing sequences remain difficult because many gliding mammals are small and nocturnal, and only a limited number of species can be induced to glide under controlled experimental conditions. Laboratory studies are also generally restricted to distances much shorter than those observed in the wild (Bishop, 2006; Essner, 2002; Paskins et al., 2007).

Several assumptions consequently limit the interpretation of this study. In the models, representative velocities reported in the literature were treated as approximations of pre-braking velocity, although some studies reported mean velocity across an entire glide. Braking was represented as a fixed proportional reduction in velocity and constant deceleration over a prescribed distance. However, it is reported that northern flying squirrels (*Glaucomys sabrinus*) do not take an aerodynamically equilibrium glide by changing their gliding velocities (Bahlman et al., 2013). The calculations did not incorporate the separate horizontal and vertical components of velocity, changes in angle of attack, patagial surface area, wing loading, or body posture. Moreover, the calculated landing energy represents translational kinetic energy immediately before contact and cannot be interpreted directly as the impact force itself, nor tissue stress.

Future models could incorporate measurable morphological parameters, including patagial surface area and aspect ratio, together with behavioral variables such as angle of attack and timing of pitch-up maneuvers. Simultaneous measurements of glide trajectory, three-dimensional velocity, braking posture, and landing force would allow the predicted energy partitioning to be tested directly.

Although the present analysis focused on mammals, gliding has evolved independently in numerous vertebrates and invertebrates (Dudley et al., 2007; Socha et al., 2015). All gliding animals must control aerial trajectories and dissipate energy before or during landing, although the relevant aerodynamic structures and landing mechanisms differ among taxa. The present framework could therefore be extended to other gliding lineages by incorporating taxon-specific relationships among distance, velocity, morphology, and braking performance data.

Overall, the present study extends the interpretation of gliding beyond the production of lift and the maximization of travel distance. It formalizes how glide distance, approach velocity, aerodynamic braking distance, and body mass jointly determine the energy remaining before contact and the deceleration required before landing. These relationships provide a quantitative basis for considering the mechanical requirements of deceleration, landing control, and energy absorption alongside conventional measures of aerial performance in studies of gliding evolution.

## Acknowledgements

The author gratefully acknowledges the researchers whose published studies and datasets formed the basis of this analysis.

## Data and resource availability

All scripts used for data processing, modeling, and analyses are available on GitHub and archived via Zenodo (DOI: 10.5281/zenodo.21619098).

## Use of AI tools

ChatGPT (OpenAI) was used to assist with the development, debugging, and revision of data analysis code, as well as to improve the clarity, grammar, and wording of the manuscript. All AI-generated suggestions and outputs were critically reviewed, verified, and revised by the author. The author takes full responsibility for the accuracy and integrity of the analysis and the final manuscript.

## Supporting Information

**Table S1. Literature-derived glide data and representative values used for distance-velocity modelling.** Reported body mass (BM), glide distance (D), and gliding velocity (V) were compiled from published studies of gliding mammals. Where ranges were reported, minimum and maximum values are shown together with representative values used for analysis. When a mean value was reported, the reported mean was used as the representative value. When only a range was available, the midpoint of the minimum and maximum values was additionally used as a representative value. When only a lower bound for glide distance was reported, the reported threshold was used as the representative distance and paired with the reported or calculated mean gliding velocity. Representative body mass was calculated as the midpoint of the ranges obtained from other literature.

**Table S2. Parameter estimates for the four candidate distance-velocity models.** Parameter estimates, standard errors (SE), and 95% confidence intervals (CI) are shown for the linear, log-distance, saturated, and saturated with *V*_0_ models fitted to the literature-derived glide data. *V*_max_, the asymptotic glide velocity; *V*_0_, the estimated glide velocity at zero distance; *L*, the characteristic distance scale of the saturated models.

**Table S3. Representative predictions from the two best-supported distance-velocity models.** Predicted gliding velocity, landing velocity, landing kinetic energy, and required deceleration are shown at glide distances of 20, 40, 60, and 80 m for the log-distance and saturated with *V*_0_ models. Landing velocity and kinetic energy were calculated assuming a 60% reduction in gliding velocity and body mass of 1 kg. Required deceleration was calculated assuming the same velocity reduction and a braking distance of 1 m.

**Table S4. Sensitivity analysis of predicted landing variables to the assumed proportional reduction in gliding velocity.** Predicted landing velocity, landing kinetic energy, and required deceleration are shown for reductions in gliding velocity of 40, 50, and 60% at glide distances of 20, 40, 60, and 80 m under the log-distance and saturated with *V*_0_ models. Landing kinetic energy was calculated assuming a body mass of 1 kg, and required deceleration was calculated assuming a braking distance of 1 m.

**Figure S1. Comparison of the four candidate models describing the relationship between glide distance and gliding velocity.** Points represent representative glide-distance and velocity values compiled from the literature (circles, *Acrobates pygmaeus*; triangles, *Galeopterus variegatus*; squares, *Glaucomys sabrinus*; crosses, *Petaurista leucogenys*; and crossed squares, *Petaurista petaurista*). Lines show predictions from the four-candidate distance-velocity models: linear (solid), log-distance (dashed), saturated (dot-dashed), and saturated with *V*_0_ (dotted).

**Figure S2. Residual diagnostics for the four-candidate distance-velocity models.** Residuals are plotted against fitted gliding velocity for the (A) linear, (B) log-distance, (C) Saturated, and (D) saturated with *V*_0_ models. Dashed horizontal lines indicate zero residual.

**Figure S3. Sensitivity analysis of predicted landing kinetic energy to body mass.** Predicted landing kinetic energy across glide distance for hypothetical body masses of 0.1, 0.5, 1.0, 1.5, and 2.0 kg under log-distance and saturated with *V*_0_ models. Predictions assume a 60% reduction in gliding velocity before landing.

**Figure S4. Sensitivity analysis of required deceleration to the assumed proportional reduction in gliding velocity.** Required deceleration across glide distance is shown for velocity reduction of 40, 50, and 60% under the log-distance and saturated with *V*_0_ models. Solid, dashed, and dotted lines indicate velocity reductions of 40, 50, and 60%, respectively. Predictions assume a braking distance of 1 m.

