## supplementary tables for "A theoretical framework for aerodynamic braking and landing in gliding mammals"

Table S1

| Reference | Species | Min. BM (g) | Max. BM (g) | Mean D (m) | Min. D (m) | Max. D (m) | Mean V (m/s) | Min. V (m/s) | Max.V (ms) | Representative BM (g) | Representative D (m) | Representative V (m/s) |
|---|---|---|---|---|---|---|---|---|---|---|---|---|
| Ando and Shiraishi, 1993 | *Petaurista leucogenys* | 1000 | 2500 | NA | 1 | 2 | NA | 3 | 4.5 | 1750 | 1.5 | 3.75 |
| Ando and Shiraishi, 1993 | *Petaurista leucogenys* | 1000 | 2500 | NA | 1 | 2 | NA | 3 | 4.5 | 1750 | 1 | 3 |
| Ando and Shiraishi, 1993 | *Petaurista leucogenys* | 1000 | 2500 | NA | 1 | 2 | NA | 3 | 4.5 | 1750 | 2 | 4.5 |
| Ando and Shiraishi, 1993 | *Petaurista leucogenys* | 1000 | 2500 | NA | 40 | 80 | NA | 8.5 | 11 | 1750 | 60 | 9.75 |
| Ando and Shiraishi, 1993 | *Petaurista leucogenys* | 1000 | 2500 | NA | 40 | 80 | NA | 8.5 | 11 | 1750 | 40 | 8.5 |
| Ando and Shiraishi, 1993 | *Petaurista leucogenys* | 1000 | 2500 | NA | 40 | 80 | NA | 8.5 | 11 | 1750 | 80 | 11 |
| Ando and Shiraishi, 1993 | *Petaurista leucogenys* | 1000 | 2500 | NA | 80 | NA | NA | 9 | 13.3 | 1750 | 80 | 11.15 |
| Bahlman et al., 2013 | *Glaucomys sabrinus* | 50 | 185 | 18 | NA | NA | 7.2 | NA | NA | 117.5 | 18 | 7.2 |
| Byrnes et al., 2008 | *Galeopterus variegatus* | 1000 | 1750 | NA | 20 | NA | 10.1 | 9.91 | 10.2 | 1375 | 20 | 10.1 |
| Krishna et al., 2016 | *Petaurista petaurista* | 1000 | 2500 | 36.3 | 7.8 | 104.3 | 8.9 | 5.5 | 13.3 | 1750 | 36.3 | 8.9 |
| Paskins et al., 2007 | *Glaucomys sabrinus* | 50 | 185 | NA | NA | NA | NA | NA | NA | 117.5 | 2.5 | 4.5 |
| Pridome and Hoffmann, 2014 | *Acrobates pygmaeus* | 12.8 | 13.6 | NA | 10 | NA | NA | NA | NA | 13.2 | 10 | 6.8 |
| Pridome and Hoffmann, 2014 | *Acrobates pygmaeus* | 12.8 | 13.6 | NA | 10 | NA | NA | NA | NA | 13.2 | 2.5 | 5.73 |
| Pridome and Hoffmann, 2014 | *Acrobates pygmaeus* | 12.8 | 13.6 | NA | 10 | NA | NA | NA | NA | 13.2 | 1 | 6.03 |
| Pridome and Hoffmann, 2014 | *Acrobates pygmaeus* | 12.8 | 13.6 | NA | 10 | NA | NA | NA | NA | 13.2 | 4 | 5.69 |
| Scheibe et al., 2006 | *Glaucomys sabrinus* | NA | NA | 14.4 | NA | NA | 6.3 | NA | NA | 125.9 | 14.4 | 6.3 |
| Scheibe et al., 2006 | *Glaucomys sabrinus* | NA | NA | 13.3 | NA | NA | 7.5 | NA | NA | 120.7 | 13.3 | 7.5 |
| Scheibe et al., 2006 | *Glaucomys sabrinus* | NA | NA | 14.1 | NA | NA | 6.4 | NA | NA | 125.4 | 14.1 | 6.4 |
| Scheibe et al., 2006 | *Glaucomys sabrinus* | NA | NA | 12.5 | NA | NA | 8.1 | NA | NA | 115.9 | 12.5 | 8.1 |
| Stafford et al., 2002 | *Petaurista leucogenys* | 1000 | 2500 | 19.46 | 4.75 | 50.35 | 6.23 | 4.39 | 9.47 | 1750 | 19.46 | 6.23 |
| Stafford et al., 2002 | *Petaurista leucogenys* | 1000 | 2500 | 19.46 | 4.75 | 50.35 | 6.23 | 4.39 | 9.47 | 1750 | 4.75 | 4.39 |
| Stafford et al., 2002 | *Petaurista leucogenys* | 1000 | 2500 | 19.46 | 4.75 | 50.35 | 6.23 | 4.39 | 9.47 | 1750 | 50.35 | 9.47 |

Table S2

| Model | Parameter | Estimate | SE | 95% CI |
|---|---|---|---|---|
| Linear | β0 | 5.25 | 0.35 | 4.51-5.98 |
| | β1 | 0.08 | 0.01 | 0.06-0.10 |
| Log-distance | α | 3.56 | 0.45 | 2.62-4.51 |
| | β | 1.49 | 0.17 | 1.14-1.84 |
| Saturated | Vmax | 8.37 | 0.47 | 7.36-9.54 |
| | L | 2.81 | 0.69 | 1.40-6.29 |
| Saturated with V0 | V0 | 4.24 | 0.47 | 3.16-5.23 |
| | Vmax | 10.88 | 0.91 | 9.34-16.63 |
| | L | 26.63 | 9.37 | 13.11-96.54 |

Table S3

| Glide distance (m) | Model | Gliding velocity (m/s) | Landing velocity (m/s) | Landing kinetic energy (J) | Required deceleration (g) |
|---|---|---|---|---|---|
| 20 | Log-distance | 8.02 | 3.21 | 5.14 | 2.75 |
| 20 | Saturated with V0 | 7.75 | 3.10 | 4.80 | 2.57 |
| 40 | Log-distance | 9.05 | 3.62 | 6.55 | 3.50 |
| 40 | Saturated with V0 | 9.40 | 3.76 | 7.07 | 3.78 |
| 60 | Log-distance | 9.65 | 3.86 | 7.46 | 3.99 |
| 60 | Saturated with V0 | 10.18 | 4.07 | 8.30 | 4.44 |
| 80 | Log-distance | 10.08 | 4.03 | 8.13 | 4.35 |
| 80 | Saturated with V0 | 10.55 | 4.22 | 8.90 | 4.77 |

Table S4

| Velocity reduction rate (%) | Glide distance (m) | Model | Landing velocity (m/s) | Landing kinetic energy (J) | Required deceleration (g) |
|---|---|---|---|---|---|
| 40 | 20 | Log-distance | 4.81 | 11.57 | 2.10 |
| 40 | 20 | Saturated with V0 | 4.65 | 10.80 | 1.96 |
| 40 | 40 | Log-distance | 5.43 | 14.73 | 2.67 |
| 40 | 40 | Saturated with V0 | 5.64 | 15.90 | 2.88 |
| 40 | 60 | Log-distance | 5.79 | 16.77 | 3.04 |
| 40 | 60 | Saturated with V0 | 6.11 | 18.67 | 3.38 |
| 40 | 80 | Log-distance | 6.05 | 18.29 | 3.31 |
| 40 | 80 | Saturated with V0 | 6.33 | 20.04 | 3.63 |
| 50 | 20 | Log-distance | 4.01 | 8.04 | 2.46 |
| 50 | 20 | Saturated with V0 | 3.87 | 7.50 | 2.29 |
| 50 | 40 | Log-distance | 4.52 | 10.23 | 3.13 |
| 50 | 40 | Saturated with V0 | 4.70 | 11.04 | 3.38 |
| 50 | 60 | Log-distance | 4.83 | 11.65 | 3.56 |
| 50 | 60 | Saturated with V0 | 5.09 | 12.96 | 3.96 |
| 50 | 80 | Log-distance | 5.04 | 12.70 | 3.88 |
| 50 | 80 | Saturated with V0 | 5.28 | 13.91 | 4.24 |
| 60 | 20 | Log-distance | 3.21 | 5.14 | 2.75 |
| 60 | 20 | Saturated with V0 | 3.10 | 4.80 | 2.57 |
| 60 | 40 | Log-distance | 3.62 | 6.55 | 3.50 |
| 60 | 40 | Saturated with V0 | 3.76 | 7.07 | 3.78 |
| 60 | 60 | Log-distance | 3.86 | 7.46 | 3.99 |
| 60 | 60 | Saturated with V0 | 4.07 | 8.30 | 4.44 |
| 60 | 80 | Log-distance | 4.03 | 8.13 | 4.35 |
| 60 | 80 | Saturated with V0 | 4.22 | 8.90 | 4.77 |
