## Supplementary figures and images for "A theoretical framework for aerodynamic braking and landing in gliding mammals"

### supplementary figures

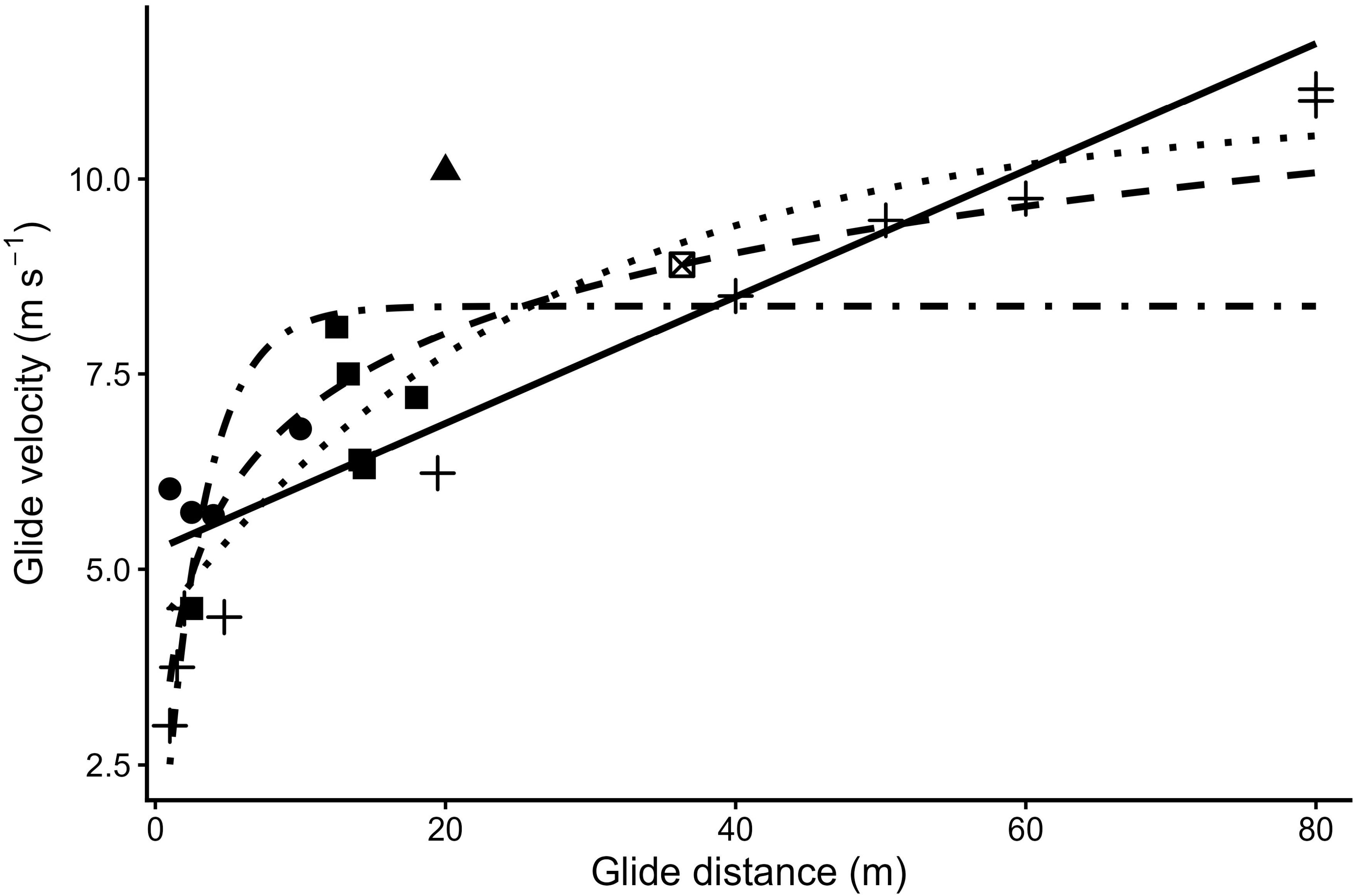

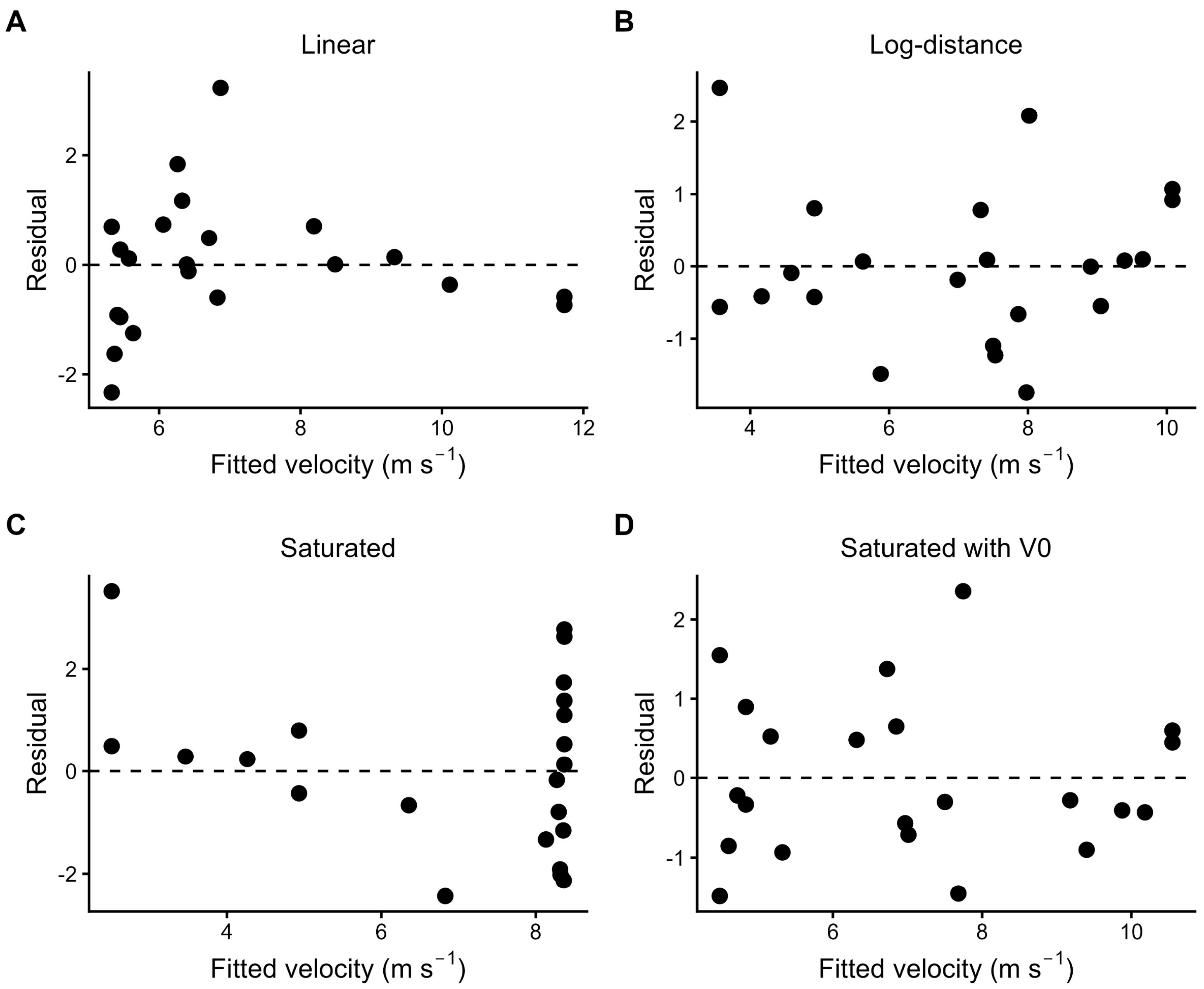

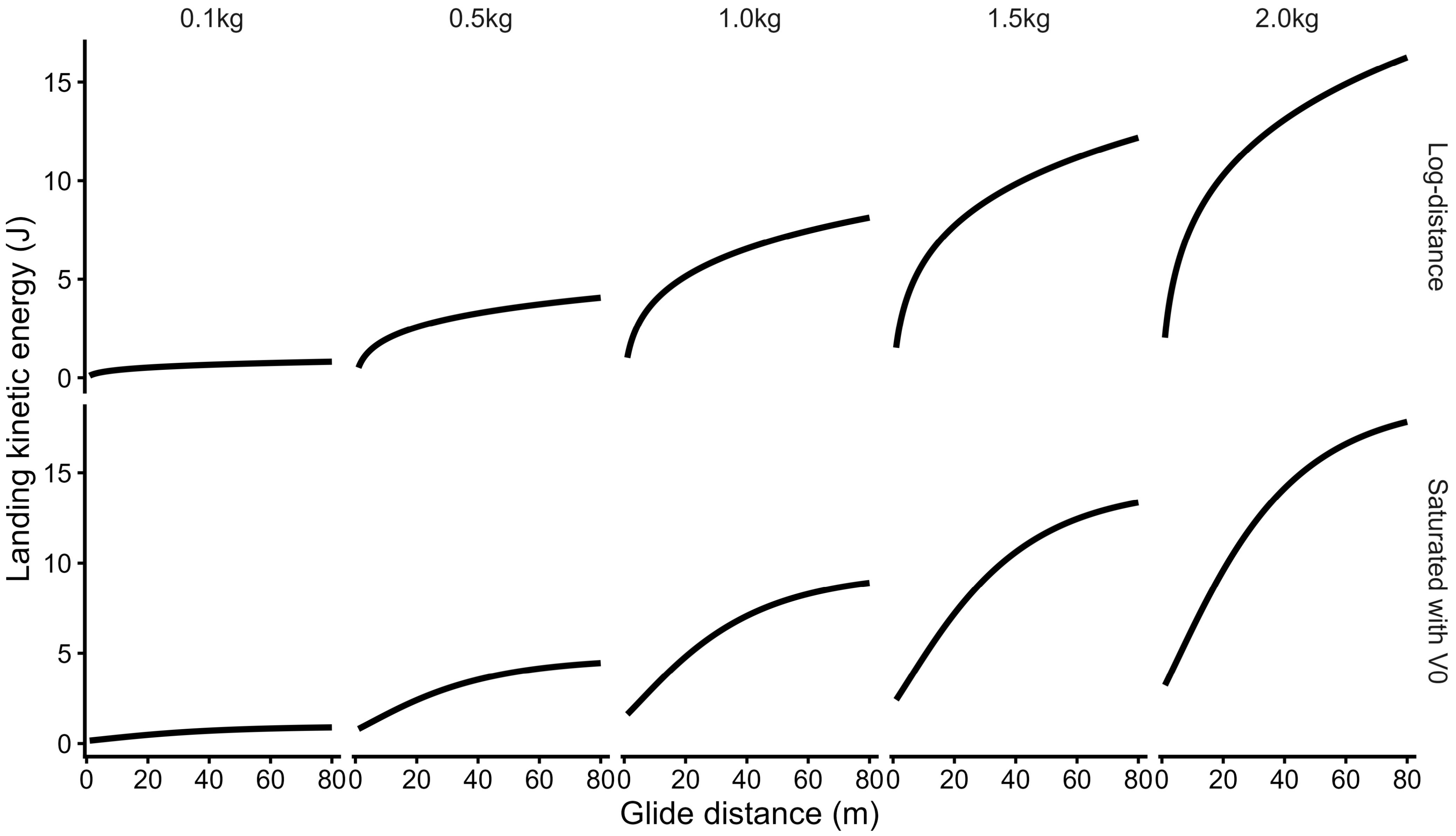

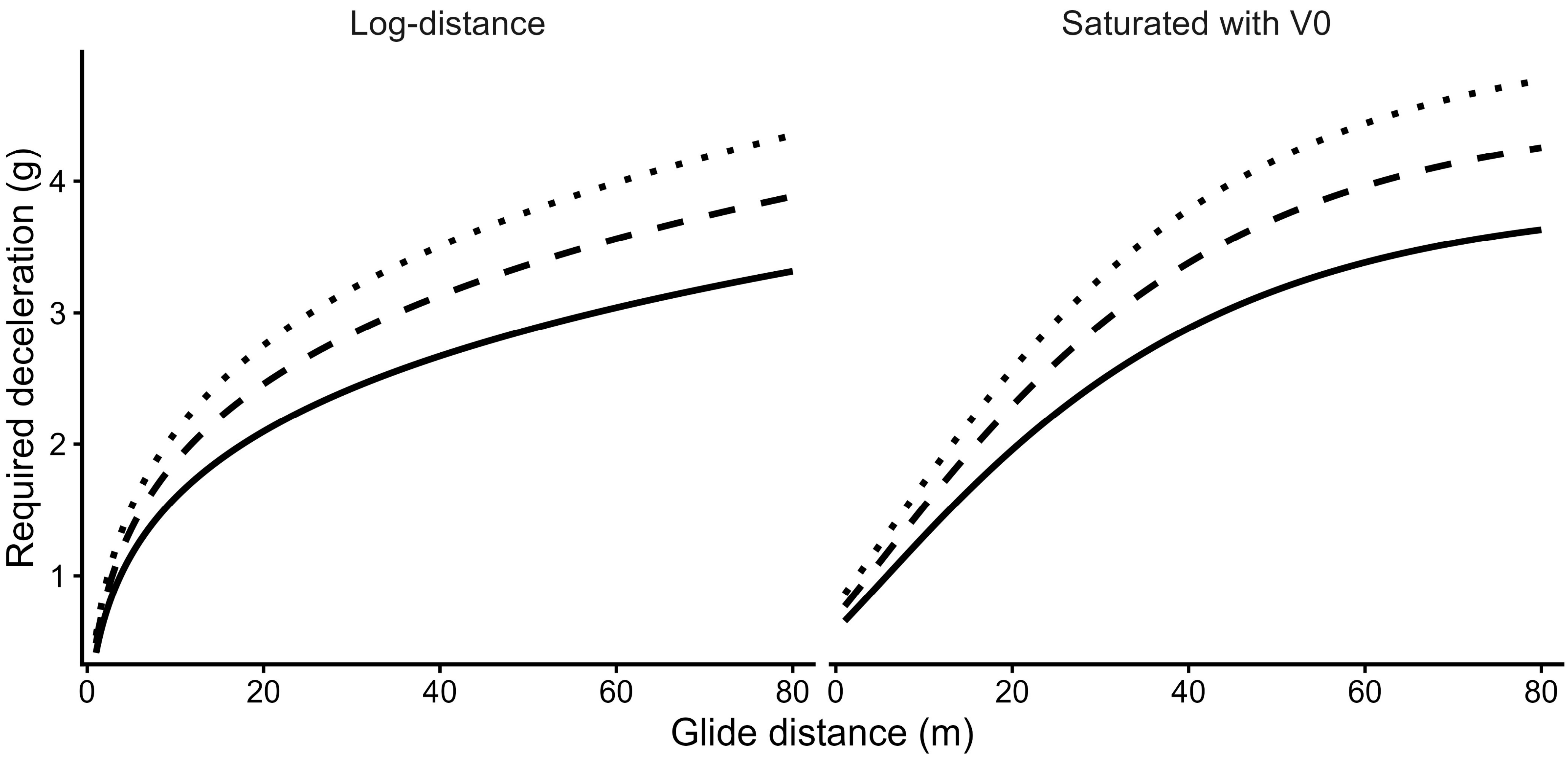
